# Isotopic Labeling Analysis using Single Cell Mass Spectrometry

**DOI:** 10.1101/2025.04.10.647921

**Authors:** Anh Hai Vu, Lorenzo Caputi, Sarah E. O’Connor

**Affiliations:** Department of Natural Product Biosynthesis, Max Planck Institute for Chemical Ecology, Jena 07745, Germany

## Abstract

Here we show that a recently developed single cell mass spectrometry method can be used to monitor the incorporation of an isotopically labeled precursor into single plant protoplasts. Specifically, we show that we can monitor the incorporation of deuterium-labeled tryptamine into an alkaloid pathway over the course of 24 hours. The resulting data provides a glimpse into the rate of synthesis and transport of chemically complex monoterpene indole alkaloids in single cells across a small population. By measuring the concentration of labeled alkaloids, we gain insight into the flux of an important plant biosynthetic pathway. Stable-isotope labeling is a widely used approach to study metabolic networks by tracking isotopologues over time. This manuscript provides proof of principle that single cell isotopic labeling can be performed in plant pathways.

## Introduction

The alkaloid biosynthetic pathways of the medicinal plant *Catharanthus roseu*s represent a particularly complex set of metabolic networks. The leaves of this plant produce dozens of biosynthetically related alkaloids including vinblastine (Fig. **S1**), an anticancer drug (O’Connor & Maresh, 2006; Zhang *et al*., 2018; Dhyani *et al*., 2022). Notably, biochemical analysis and single cell RNA-sequencing (scRNA-seq) have shown that the complexity of this system extends to spatial organization: the biosynthetic genes of these alkaloid pathways are partitioned among three cell types (Courdavault *et al*., 2014; Kulagina *et al*., 2022; Li *et al*., 2023; Sun *et al*., 2023). Moreover, a recently developed single cell mass spectrometry (scMS) method has provided insight into the location and concentrations of the biosynthetic intermediates and products in specific leaf cells (Vu *et al*., 2024). Specifically, the early genes of the pathway are found in Internal Phloem Associated Parenchymal (IPAP) cells (Guirimand, G. *et al*., 2011; Li *et al*., 2023; Sun *et al*., 2023). The intermediate loganic acid (Fig. **S1**) is transported from the IPAP cells to epidermal cells, where further biosynthetic steps take place (Guirimand, Grégory *et al*., 2011; Li *et al*., 2023; Sun *et al*., 2023). scMS has shown that secologanin (Fig. **S1**), a downstream product of loganic acid, accumulates to levels of up to 500 mM in individual cells (Vu *et al*., 2024). Secologanin serves as a precursor for the epidermal-localized enzyme strictosidine synthase (STR), which catalyzes the reaction of secologanin with tryptamine to form strictosidine (Fig. **S1**). Strictosidine is subsequently converted by downstream epidermal-localized enzymes to assemble a variety of alkaloids including ajmalicine, stemmadenine acetate, tabersonine and catharanthine (Fig. **S1**). Late-stage alkaloids are then exported to idioblast cells where some are further derivatized by idioblast-localized enzymes (Guirimand, G. *et al*., 2011; Li *et al*., 2023; Sun *et al*., 2023) (Fig. **1a**). scMS shows that late-stage alkaloids catharanthine and vindoline accumulate to high levels (concentrations up to 100 and 50 mM, respectively) in individual cells (Yamamoto *et al*., 2019; Li *et al*., 2023; Vu *et al*., 2024). Vindoline and catharanthine are ultimately dimerized and derivatized to form anhydrovinblastine and vinblastine (Fig. **S1**) (Goodbody *et al*., 1988).

**Fig 1.**
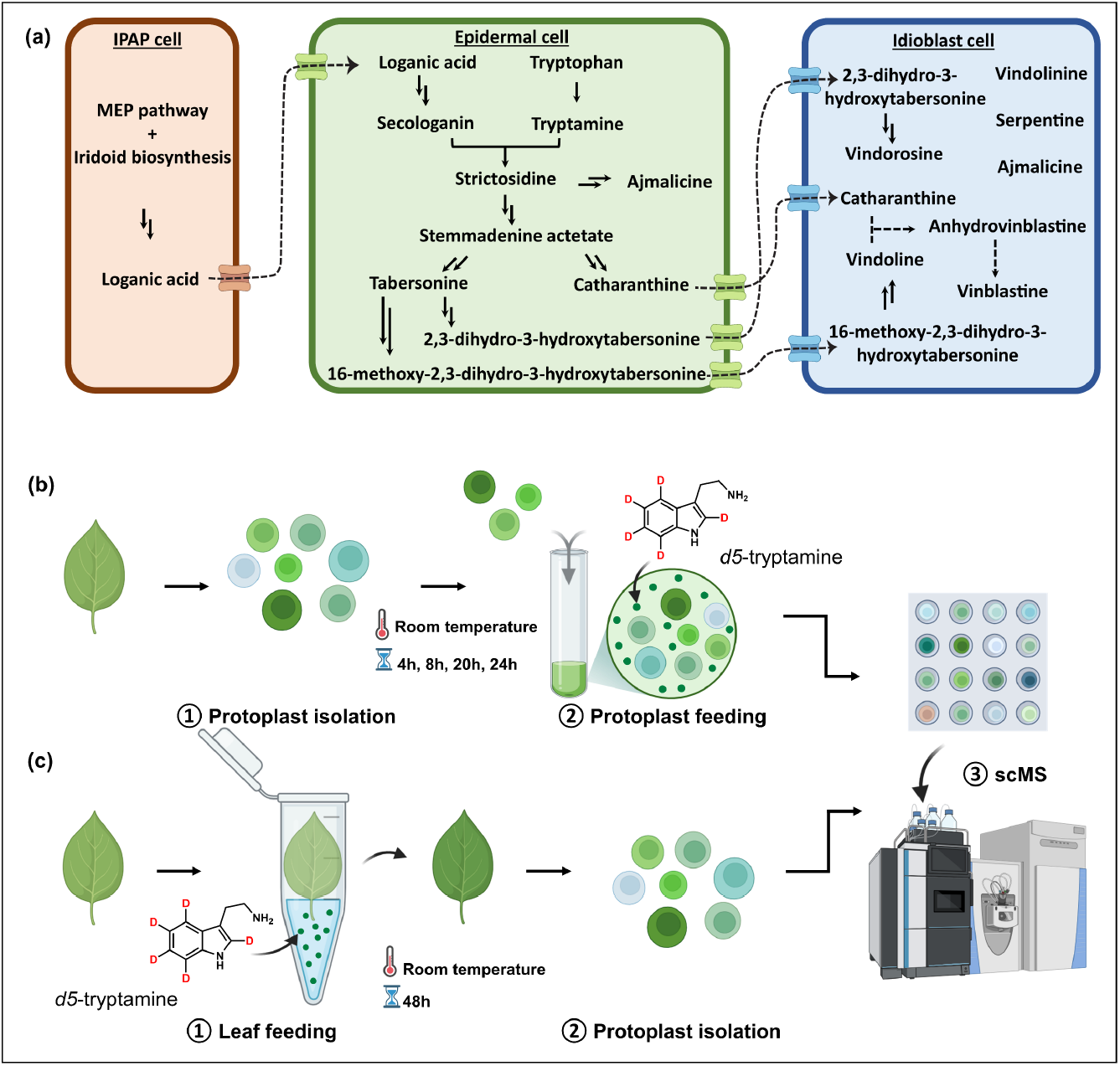
(a) A simplified schematic of the MIA biosynthetic pathway. The chemical structures of the compounds are shown in Fig. **S1**. Workflow for two scMS feeding experiments is shown. (b) Isotopically labeled substrate is fed directly to protoplast suspension in MM buffer (0.4 M mannitol, 20 mM MES, pH 5.7), incubated at different time points, and then analyzed by scMS. (c) Alternatively, isotopically labeled substrate is fed through the petiole of detached leaves, followed by protoplast isolation from the fed leaves and analysis by scMS.

Stable-isotope labeling– the feeding of an isotopically labeled precursor to a whole cell or organism– is a widely used approach to establish the starting materials and intermediates of a metabolic pathway and to obtain a dynamic model of the metabolic networks by tracking isotopologues over time (Chokkathukalam *et al*., 2014; Doppler *et al*., 2019; Fernández-García *et al*., 2020; Arroo *et al*., 2021; Yu *et al*., 2023). Here we use scMS to quantitatively monitor the incorporation of the isotopically labeled precursor *d5*-tryptamine (Fig. **S1**) into the *C. roseus* alkaloid pathway at the single cell level. We observed the formation of labeled alkaloids *d4*-strictosidine, *d4*-ajmalicine, *d4*-stemmadenine acetate, *d4*-catharanthine and *d4*-tabersonine (Fig. **S1**) in individual protoplasts. Quantification of the *d4*-strictosidine levels over this 24-hour time frame allowed us to calculate the rate of enzymatic formation of this compound per cell. *d5*-tryptamine was also incubated with an intact *C. roseus* leaf that was then analyzed by scMS after 48 hours (Fig. **1c**). This analysis revealed a profile of deuterated alkaloids distinct from those produced in the protoplast solution, suggesting that the leaf structure impacts the fidelity of this biosynthetic pathway. Overall, this study demonstrates that scMS can be used to track the isotopic labeling of complex natural products in single plant cells.

## Results and Discussion

### Metabolic capacity of C. roseus protoplasts in bulks

We first assessed whether protoplasts were metabolically active. Protoplasts were generated from *C. roseus* leaf tissue as reported previously (Carqueijeiro *et al*., 2018; Vu *et al*., 2024) and incubated with *d5*-tryptamine. Aliquots of protoplasts were harvested at specific time points and the incorporation of deuterium into the alkaloids was evaluated by standard LC-MS. We observed formation of *d4*-strictosidine, *d4*-ajmalicine, *d4*-stemmadenine acetate, *d4*-catharanthine, *d4*-tabersonine (Fig. **S2**). Formation of *d4*-strictosidine was observed in the first time point (less than 5 minutes after the addition of *d5*-tryptamine) but the later intermediate *d4*-stemmadenine acetate could be detected only after 2.5 h. Downstream products *d4*-catharanthine and *d4*-tabersonine were first observed at the 6 h time point. The levels of labeled products appeared to plateau at 40 hours, suggesting that the metabolic activity of the protoplasts had slowed or stopped by this time point. Cell viability was monitored over time during feeding (Fig. **S3**). After 24 hours, the cell viability of protoplasts incubated with 1 mM *d5*-tryptamine was 50%, whereas only 20% of cells incubated with 5 mM *d5*-tryptamine remained viable. For this reason, protoplast feeding for single cell analysis was performed with 1 mM *d5*-tryptamine over 24 hours. While it is likely that protoplasts export alkaloids into the media, it would be impossible to determine whether alkaloids in the media came from specific export processes, or are simply derived from unviable protoplasts that lysed during the experiment. Therefore, the alkaloid profile of the media was not considered in these measurements.

### scMS of isotopically labeled protoplasts

Using the conditions optimized for bulk protoplasts described above, we monitored the formation of *d5*-tryptamine derived isotopologues in single cells over time (Fig. **1b**). Aliquots of leaf protoplasts incubated with 1 mM *d5*-tryptamine were taken at 4 h, 8 h, 20 h and 24 h, loaded onto a SieveWell™ chip, picked and then subjected to scMS analysis to quantify the levels of both natural compounds and the corresponding isotopologues. Similar to the mass spectrometry profile observed with bulk solutions of protoplasts, we observed the formation of *d4*-strictosidine at the first time point (4 h), and formation of *d4*-ajmalicine at the 8 h time point. *d4*-Stemmadenine acetate, *d4*-catharanthine and *d4*-tabersonine were first observed at the 20 h time point (Figs **2a-c**). These observations clearly demonstrate that it is possible to monitor stable isotope incorporation in single cells using mass spectrometry.

**Fig 2.**
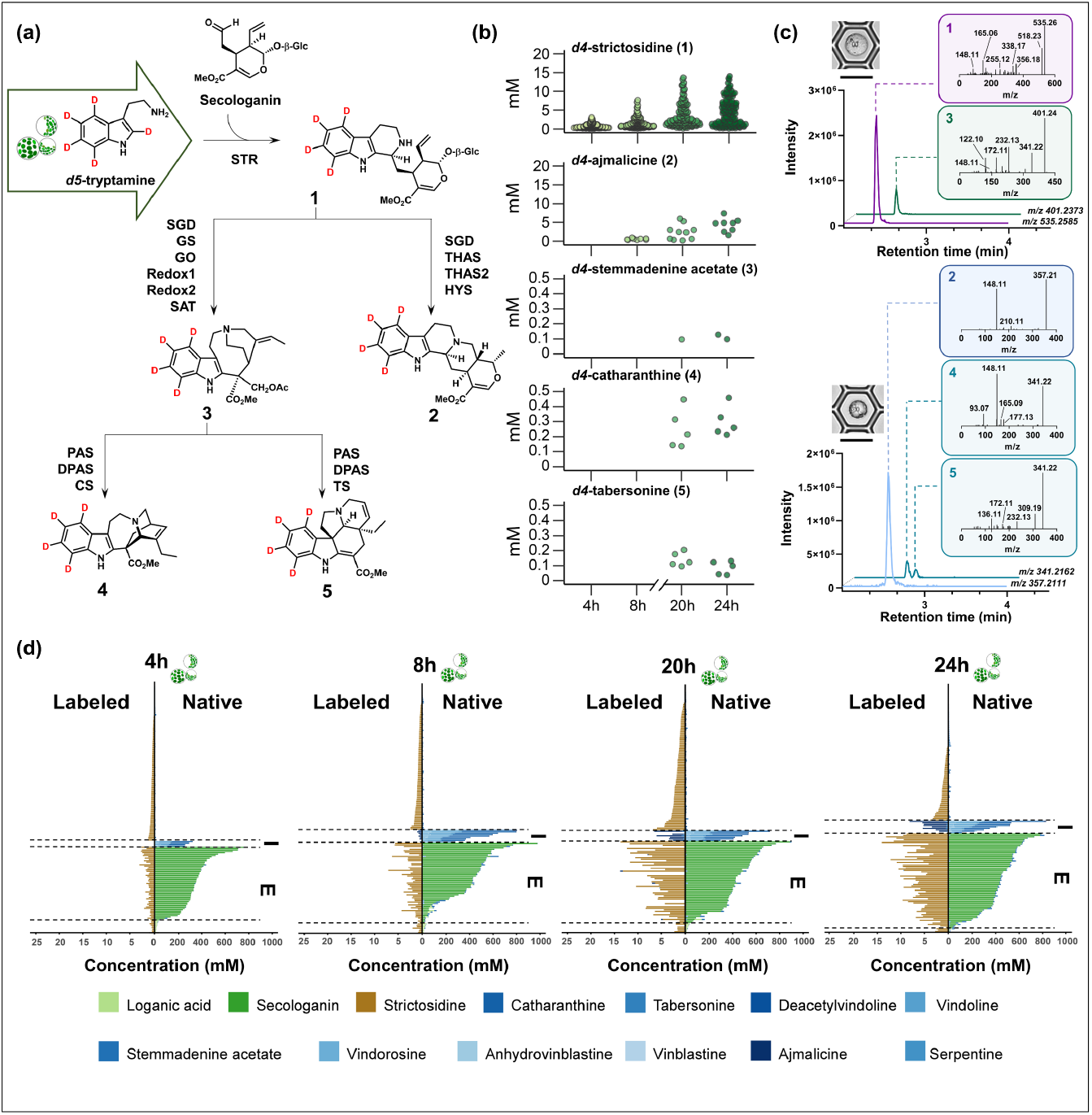
Metabolic profile of single cells from protoplast-feeding experiments. (a) Biosynthesis scheme of labeled metabolites detected. (b) Calculated intracellular concentration of detected labeled products in a single cell over time. (c) Representative cells and the corresponding chromatograms and MS/MS fragmentation of labeled products. (d) Stacked bar chart showing native and labeled metabolic profiles of single cells at four time points. A wide variety of endogenous, unlabeled compounds were also observed in this cell population. Chemical structures for all compounds are shown in Fig. S1.

We used two unlabeled compounds, secologanin and vindoline, as molecular markers to identify epidermal and idioblast cells, respectively, in the analyzed cell population. *d4*-Ajmalicine, *d4*-catharanthine and *d4*-tabersonine were observed to be present in vindoline-containing idioblast cells (Fig. **2d**). Based on the location of the biosynthetic genes, these three compounds are expected to be synthesized in epidermal cells (Li *et al*., 2023; Sun *et al*., 2023), so this observation suggests that these compounds are exported from epidermal protoplasts, and then selectively imported into idioblast protoplasts. *d4*-Stemmadenine acetate, which is a biosynthetic intermediate on pathway to tabersonine and catharanthine (Fig. **1a**), was found primarily in secologanin containing epidermal protoplasts (Fig. **S4**), suggesting that this compound is retained within epidermal cells. No other known alkaloid biosynthetic intermediates were detected, either because these intermediates are unstable or because they are consumed *en route* to one of the downstream alkaloid products.

Strikingly, after 4 h, *d4*-strictosidine was detected in ca. 85% of the analyzed cells (Fig. **S4**). However, only 30% of the analyzed cells were shown to contain secologanin, which is the immediate precursor of strictosidine (Fig. **1a**). Therefore, secologanin containing cells (likely epidermal cells) must produce *d4*-strictosidine and then export this compound into the medium, where it is imported into a large fraction of cells. This observation suggests that many cells have the capacity to import strictosidine. Notably, idioblasts (assigned by the presence of unlabeled vindoline) do not take up *d4*-strictosidine, highlighting the selectivity of the idioblast import processes (Fig. **2d**). Although strictosidine could form non-enzymatically from secologanin and *d5*-tryptamine, this non-enzymatic reaction would also produce the non-natural stereoisomer, vincoside (*R*-isomer) (Fig. **S5**), which was never detected in our experiments. Therefore, we assume that all observed *d4*-strictosidine was formed enzymatically.

### Calculating reaction rate

We attempted to calculate the rate of synthesis of these labeled compounds per cell, though the active transport of these labeled compounds from the cell type of synthesis complicated this calculation. scRNA-seq data indicates that only epidermal cells express the biosynthetic genes required for production of *d4*-strictosidine, *d4*-ajmalicine, *d4*-catharanthine and *d4*-tabersonine. Epidermal cells represent 30-40 % of the total number of protoplasts analyzed in these scMS experiments, as evidenced by the presence of the molecular marker secologanin (Table **S3**, Supporting Information Dataset **S1**). Therefore, the total amount of labeled compounds accumulated at each time point, expressed in femtomoles (fmol), was calculated and divided by the number of epidermal (secologanin-containing) protoplasts. For instance, the total amount of *d4*-strictosidine produced after 4 h of feeding was 1130 fmol. This amount was divided by the number of cells containing secologanin (epidermal cells) to obtain an estimate of the absolute amount of *d4*-strictosidine produced by a single cell in 4 h (Table **S3**, Dataset **S1**). This value increased over time to 64.86 fmol/cell after 20 h. No further increase was observed between 20 and 24 h. The highest intra-cellular concentration of *d4*-strictosidine measured after 24 h was 14 mM (Fig. **2b**). We note that this calculated rate may be impacted by the high levels of the supplemented *d5*-tryptamine substrate (1 mM).

The relatively simple heteroyohimbine alkaloid *d4*-ajmalicine, which is formed by the action of two enzymes (SGD, HYS) on *d4*-strictosidine (Fig. **2a**), was observed after 8 h of incubation only in a small number of cells; these cells were assigned as idioblasts based on the presence of vindoline. The amount of *d4*-ajmalicine increased significantly over time from 4.15 fmol after 8 h to 37.91 fmol after 24 h, corresponding to average intracellular concentrations of 0.56 and 7.41 mM, respectively (Fig. **2b**). Isotoplogues of the more complex alkaloids catharanthine and tabersonine were first observed after 20 hours. The average amount of *d4*-catharanthine was 2.15 and 3.17 fmol after 20 h and 24 h, respectively; whilst *d4*-tabersonine was 1.07 and 1.27 fmol at the same time points (Dataset **S1**).

### Isotopic labeling in intact leaf

We also incubated an intact, detached leaf into a solution of labeled substrate to evaluate the rate of formation of isotopologues in the cells when feeding is performed on an intact organ. Initial experiments performed in the same conditions as the protoplast feeding (1 mM *d5*-tryptamine for 24 h) did not lead to formation of detectable amounts of isotopologues when bulk leaf tissue was analyzed (Fig. **S6**). However, when 5 mM *d5*-tryptamine and a 48-hour incubation time were used, labeled products were detected, so these conditions were used. After feeding, we dissociated the tissue to release the protoplasts, which were then analyzed by scMS (Fig. **1c**). Under these conditions, only a small percentage of the analyzed cells accumulated *d4*-strictosidine (11%); approximately half of these cells contained secologanin, suggesting that in intact leaf, *d4*-strictosidine was more likely to be retained in the epidermal cells (Fig. **S7**). The average accumulation of *d4*-strictosidine was 8.19 fmol/cell and the highest intra-cellular concentration observed was 4.05 mM, which is significantly lower than the levels observed when *d5*-tryptamine was fed to protoplasts directly (Fig. **S8, 2d**, Dataset **S1**). In addition to *d4*-strictosidine, other isotopologues, *d4*-ajmalicine, *d4*-catharanthine and *d4*-tabersonine, were detected in the single cells at concentrations comparable to those observed in the protoplast feeding experiment. In contrast to experiments using detached protoplasts, we could detect accumulation of isotopologues of the more highly derivatized alkaloids vindoline and vindorosine. The highest amounts detected in a single cell were 72.32 fmol (*d4*-vindorosine) and 36.54 fmol (*d3*-vindoline) (Fig. **S1**, Dataset **S1**). Gene localization data suggests that compounds are formed after transport of 16-methoxy-2,3-dihydro-3-hydroxytabersonine and 2,3-dihydro-3-hydroxytabersonine from epidermal to idioblast cells, at which point, the idioblast-localized enzymes NMT, D4H and DAT can convert these intermediates into vindoline and vindorosine, respectively (St-Pierre *et al*., 1999; Qu *et al*., 2015; Liu *et al*., 2021; Kulagina *et al*., 2022) (Figs **1a, S1**). No labeled anhydrovinblastine or vinblastine were observed in any of these experiments, indicating that these late-stage compounds were not synthesized from labeled tryptamine under these conditions.

## Conclusion

Combining stable isotope tracing with single cell mass spectrometry provides a high-resolution window into metabolic processes across a population of cells. Only a few studies that combine isotopic labeling with single cell mass spectrometry have been reported. These studies focused on mammalian and microbial systems, and moreover, used MALDI-MS or nanoSIMS, which are mass spectrometry approaches that cannot be used to accurately quantify the absolute levels of cellular metabolites (Alcolombri *et al*., 2022; Wang, G *et al*., 2022; Wang, L *et al*., 2022; Buglakova *et al*., 2024; Xiang *et al*., 2024; Rietjens & Heijs, 2025). Here we use a recently reported scMS method to quantitatively track conversion of *d5*-tryptamine into downstream alkaloid metabolites in individual plant protoplasts. This study highlights that we can track the synthesis and transport of plant natural products at single cell resolution in real time.

## Materials and methods

### Plant growth conditions

*Catharanthus roseus* (*C. roseus*) (L.) G. Don. (Sunstorm Apricot cultivar) plants were germinated and grown in a York chamber at 23 °C, under a 16h/8h light/dark cycle.

### Chemicals and reagents

The following chemicals were used as analytical standards: L-tryptophan-(*indole-d5*) (Sigma Aldrich), secologanin (Sigma Aldrich), ajmalicine (Sigma Aldrich), catharanthine (Abcam), deacetylvindoline (Toronto Research Chemicals Inc.), anhydrovinblastine disulfate (Toronto Research Chemicals Inc.), vinblastine sulfate (Thermo Scientific Chemicals), serpentine hydrogen tartrate (Sequoia Research Products Ltd.) loganic acid (Extrasynthese) and ajmaline (Extrasynthese), tabersonine (Tokyo Chemical Industry Co. Ltd.), vindolinine (Advanced ChemBlocks Inc.) and vindoline (Acros Organics). Tryptamine-(*indole-d5*) (*d5*-tryptamine) and stemmadenine acetate were synthesized as previously reported (Kamileen *et al*., 2022; Lombe *et al*., 2025). Strictosidine was prepared as described by Boccia *et al*. (Boccia *et al*., 2022). Vindorosine was purified by reverse phase HPLC from a preparation of vindoline in which it appeared as a contaminant.

For protoplast extraction, cellulase Onozuka R-10, macerozyme R-10 were obtained from SERVA; pectinase, mannitol, KCl, and MES from Sigma Aldrich. All solvents used in this study were UHPLC/MS grade.

### Enzymatic synthesis of *d5*-tryptamine from *d5*-tryptophan

*d5*-tryptamine was synthesized by decarboxylation of L-tryptophan-(*indole-d5*), using *Ruminococcus gnavus* tryptophan decarboxylase (RgnTDC) (McDonald *et al*., 2019). The reaction mix (total volume 10 mL) contained L-tryptophan-(*indole-d5*) (10 mM), pyridoxal 5′-phosphate (PLP, 100 μM), and *Rgn*TDC (200 nM) in potassium phosphate buffer (pH 8.0, 50 mM). The reaction was incubated at 37 °C for 20 h and was stopped by adding 10 mL acetonitrile (ACN). The mixture was centrifuged at 20,000 x *g* for 10 minutes to precipitate the enzyme. The pellet was discarded and ACN was evaporated using a rotary evaporator. The aqueous suspension was purified through acid-base extraction. The pH was adjusted to approximately 12-13 by dropwise addition of 10 M NaOH. The basic layer was washed 3 times with 10 mL ethyl acetate (EtOAc). The organic layers were combined and then concentrated on a rotary evaporator to a volume of 10 mL. The EtOAc layer was further extracted 3 times with 10 mL HCl (10 mM). All aqueous phases were collected, snap-frozen, and lyophilized to yield purified *d5*-tryptamine hydrochloride powder. Product purity was verified by LC/MS and structural characterization was performed by comparing the MS/MS fragmentation and retention time of the obtained isotope labeled product to the spectrum and retention of an unlabeled authentic standard.

### Protoplast isolation

The protoplast isolation process was described in detail in a previous publication (Vu *et al*., 2024). Briefly, 1 g of young healthy leaves (2-3 cm long) was harvested and digested for 2 h at room temperature in an enzyme solution (2% cellulase R-10, 0.3% macerozyme R-10, 0.1% pectinase) prepared in MM buffer (0.4 M mannitol and 20 mM MES, pH 5.8). After passing through 70- and 40-μm strainers, protoplasts were centrifuged at 70 x *g* for 5 min to remove the supernatant containing the enzyme. Cells were washed with MM buffer, followed by centrifugation a total of three times.

### Incubation of *d5*-tryptamine with *C. roseus* leaf protoplasts: mass spectrometry analysis in bulk

After preparation as described above, protoplasts (approx. 20 mL) were aliquoted in 48-well plates and assigned to different treatment groups. Protoplasts were incubated with either 0 mM, 1 mM, 5 mM *d5*-tryptamine prepared in MM buffer, and 600 μL aliquots were taken at specified time points (less than 5 minutes, 15 minutes, 30 minutes, 1 hour, 1.5 hours, 2 hours, 2.5 hours, 3 hours, 4 hours, 5 hours, 6 hours, 7 hours, 8 hours, 16 hours, 20 hours, 24 hours, 40 hours, 48 hours, 53 hours post incubation). The collected protoplast suspension was homogenized and 2.5 μL of this homogenized solution was extracted with 100 μL aqueous methanol (50/50, v/v), with 10 nM ajmaline as the internal standard. The extract was then filtered and subjected to LC-MS analysis to monitor the conversion of *d5*-tryptamine into downstream alkaloids. Cell viability at each time point was measured by counting the protoplasts stained with fluoresceine diacetate divided by the total protoplast counted under bright field on a hemocytometer. Counts were performed in triplicate.

### Incubation of *d5*-tryptamine with *C. roseus* leaf protoplasts: isolation of single protoplasts for mass spectrometry analysis

After protoplast isolation, aliquots of 200 μL of cell suspension were added into 2 mL Eppendorf tubes. To start the incubation, 200 μL of 2 mM *d5*-tryptamine was added to each tube. Each tube served as a sample for a time point for scMS analysis (4 h, 8 h, 20 h, 24 h). Single cell picking was performed as previously described (Vu *et al*., 2024). Briefly, approximately 10,000 protoplasts were distributed and trapped onto a microwell membrane with 50 μm micropore size and imaged by bright-field microscopy to record size and morphology. Single cells were picked using a microfluidics-based robot and transferred to a 96-well plate containing 8 μL of 0.1% formic acid in water. Next, 8 μL of methanol containing internal standard was added to disrupt the cells completely and release the metabolites. LCMS analysis was then performed as described below. The total number of cells analyzed is shown in Table **S1**.

### Incubation of *d5*-tryptamine with *C. roseus* leaf: mass spectrometry analysis of single protoplasts

Six 2–3-cm long *C. roseus* leaves were placed in a solution of 100 μL of either 1 mM or 5 mM *d5*-tryptamine, where the labeled precursor was absorbed into the leaf though the cut petioles. The leaves were collected for protoplast isolation and scMS analysis 24 and 48 hours after the start of the experiment. An additional 100 μL aliquot of water was added to the leaves after 24 hours. Leaves were then collected for bulk MS analysis, protoplast isolation and scMS analysis.

### LC-MS analysis of *d5*-tryptamine-derived metabolites

LC-MS analysis was carried out as described previously (Vu *et al*., 2024). In brief, UHPLC-HRMS analysis was performed on a Vanquish (Thermo Fisher Scientific) system coupled to a Q-Exactive Plus (Thermo Fisher Scientific) Orbitrap mass spectrometer. For LC separation, a Waters™ ACQUITY UPLC BEH C18 130 Å column (1.7 μm, 2.1 mm x 50 mm) column was used at a temperature of 40°C. The binary mobile phases consisted of MilliQ water with 0.1 % formic acid (A) and acetonitrile (B). The 7-minute gradient started with a linear increase from 1% to 70% B for 5 min. The wash stage was set at 99% B for 0.5 min before switching back to 1% B for 1.5 min to condition the column for the next injection. The flow rate was 0.3 mL min^-1^, the injection volume was 4 μL, and the sample tray was kept at 10 °C.

Mass spectrometry data acquisition was performed in full scan MS mode, at resolution 70,000 in positive mode over the mass range *m/z* from 120 to 1,000. The full-scan and data-dependent MS/MS mode (full MS/dd-MS^2^ Top10) were used for QC pooled samples to simultaneously record the spectra of the precursors and fragmentation. In addition, the full MS/dd-MS^2^ mode with an inclusion list of targeted compounds was applied to the pooled QC samples to confirm fragments of the selected precursors. The parameters for dd-MS^2^ were set as follows: resolution 17,500, mass isolation window 0.7 Da, and normalized collision energy (NCE) was set at 3 levels: 15%, 30%, and 45%. The spectrum data format was centroid. All the parameters of the UHPLC-HRMS system were controlled through Xcalibur software version 4.3.73.11 (Thermo Fisher Scientific). Chromatographic peak areas from extracted ion chromatograms (EIC) were integrated and extracted using the Xcalibur Quan Browser version 4.3.73.11 (Thermo Fisher Scientific).

### Identification and quantification of targeted compounds in a single cell

The identification of targeted compounds was confirmed by comparing retention times and MS/MS fragmentation patterns to those of authentic standards. Standard solutions were prepared in methanol (MeOH) at approximately 1 mM concentration (exact concentration was recorded). Serial dilutions were performed down to 0.001 nM, and these solutions were analyzed by UHPLC-MS to estimate the limit of quantification (LOQ) and the calibration range (Table **S2**). Each calibration point was measured in triplicate, and linear regression curves were generated using peak areas. Deuterium labeled compounds were quantified using calibration curves of the corresponding unlabeled standards. The absolute amount of analyte in each single cell (fmol) was determined using the external calibration curve. Analyte concentration (mM) in a single cell was calculated by dividing the absolute amount of analyte (fmol) by the estimated cell volume, assuming the cells are spherical. The diameter of each cell was measured using ImageJ software based on cell images recorded by the cell-picking robot. Data from wells containing no cell or more than one cell due to unsuccessful cell picking were excluded.

### Data analysis

Statistical analyses were performed using R (version 4.3.1). Violin point graphs and dot plots were visualized by the ggplot2 (version 3.5.0) package. Stacked bar charts were visualized by GraphPad Prism software (version 9.5.1).

## Author Contributions

AHV conducted all experiments. AHV and LC performed data analysis. AHV, LC, SEO conceived of the project and wrote the manuscript.

## Acknowledgements

We thank Dr. Mohamed Omar Kamileen, Dr. Allwin McDonald, Dr. Gyumin Kang, Clara Morweiser, and Abdullah Sandhu for providing authentic standards. We would like to thank Sarah Heinicke and Dr. Maritta Kunert for their assistance with mass spectrometry analysis, and Jens Wurlitzer for his support with the cell-picking robot.

