## Supplementary Figures for "Isotopic Labeling Analysis using Single Cell Mass Spectrometry"

### Table of Contents

**Pages**

**Supplementary Figures** S3 – S13

**Fig. S1** Compound structures and names mentioned in this work.

**Fig. S2** Incorporation of *d5*-tryptamine in bulk leaf protoplasts derived from *C. roseus* leaf.

**Fig. S3** Analysis of cell viability under three different feeding conditions: 0 mM, 1 mM, and 5 mM of *d5*-tryptamine in MM Buffer (0.4 M mannitol, 20 mM MES, pH 5.7).

**Fig. S4** Metabolic profile of both labeled and native target compounds quantified in single protoplasts over a 24-hour time course of the protoplast feeding experiments

**Fig. S5** Extracted ion chromatograms (EICs) and MS/MS spectra of strictosidine and vincoside standards.

**Fig. S6** Single cell metabolic profile of leaves fed through the petiole with 1 mM *d5*-tryptamine for 24 hours.

**Fig. S7** Single cell metabolic profile of leaves fed through the petiole with 5 mM *d5*-tryptamine for 48 hours.

**Fig. S8** Intracellular concentration in mM of labeled products in a single cell over time in the intact leaf feeding experiments

### Supplementary Tables S14 – S15

**Table S1** Number of cells analyzed in scMS experiments.

**Table S2** Analytical parameters of the compounds quantified in this study.

**Table S3** Data used to calculate the rate of accumulation of *d4*-strictosidine in the protoplast feeding experiment.

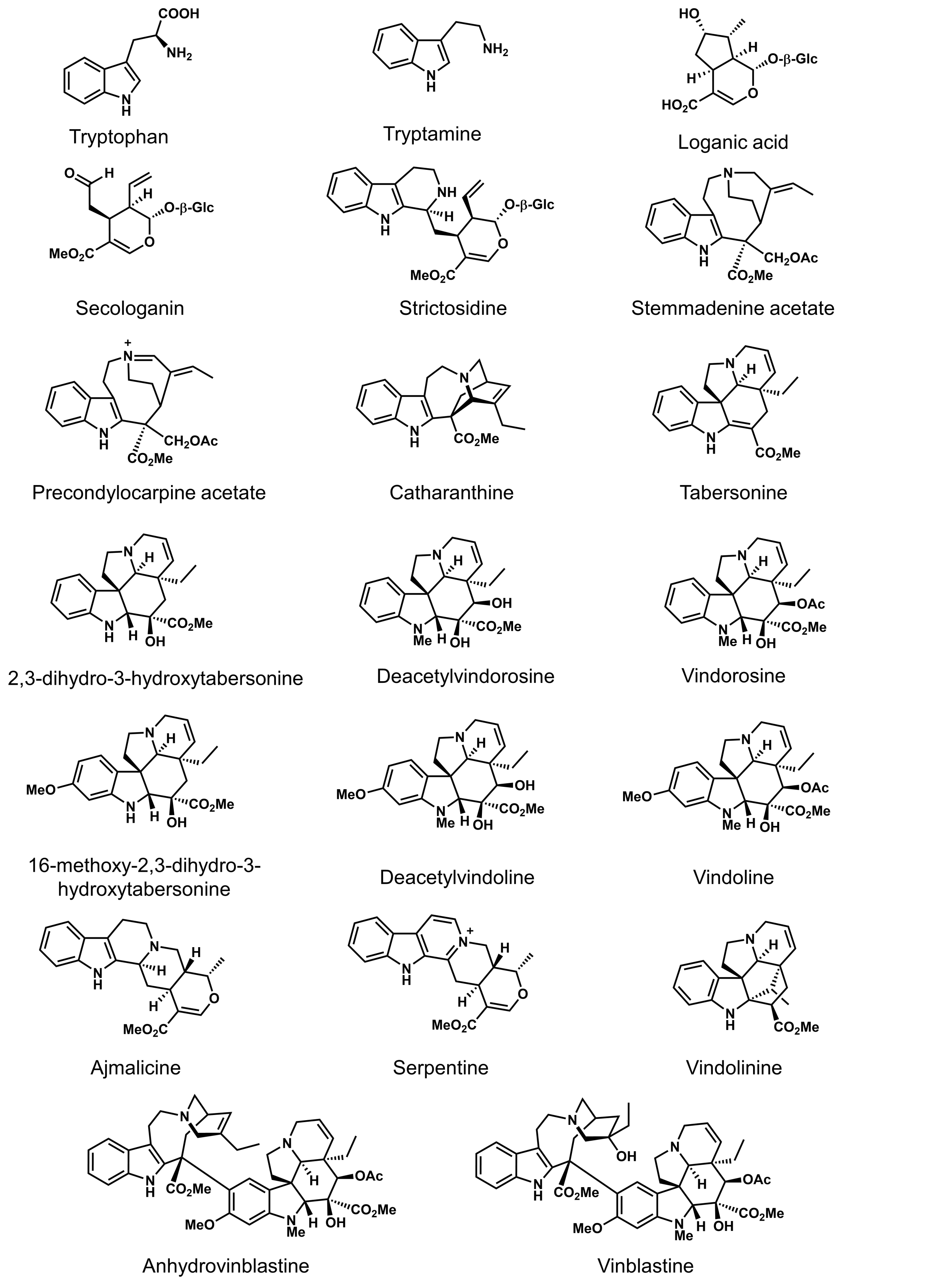


**Figure S1**

Continued on next page →

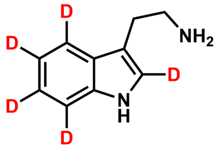

*d5-t*ryptamine

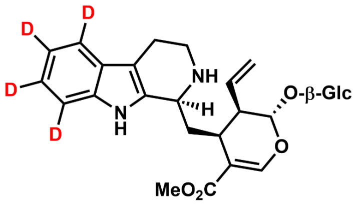

*d4*-strictosidine

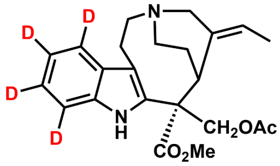

*d4*-stemmadenine acetate

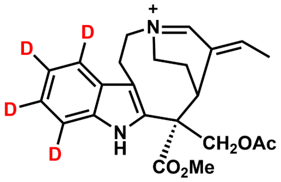

*d4*-precondylocarpine acetate

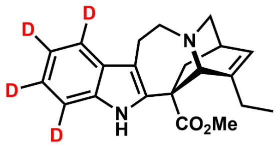

*d4*-catharanthine

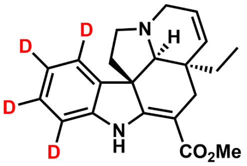

*d4*-tabersonine

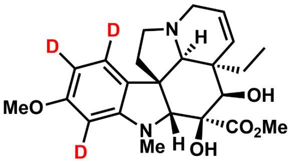

*d3*-deacetylvindoline

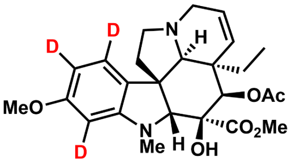

*d3*-vindoline

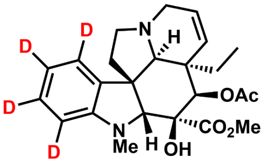

*d4*-vindorosine

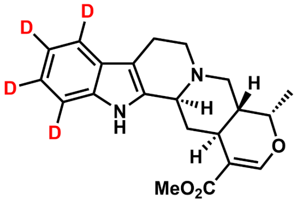

*d4*-ajmalicine

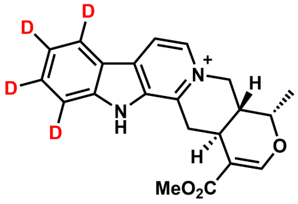

*d4*-serpentine

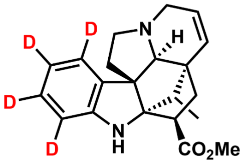

*d4*-vindolinine

**Figure S1.** Compound structures and names mentioned in this work.

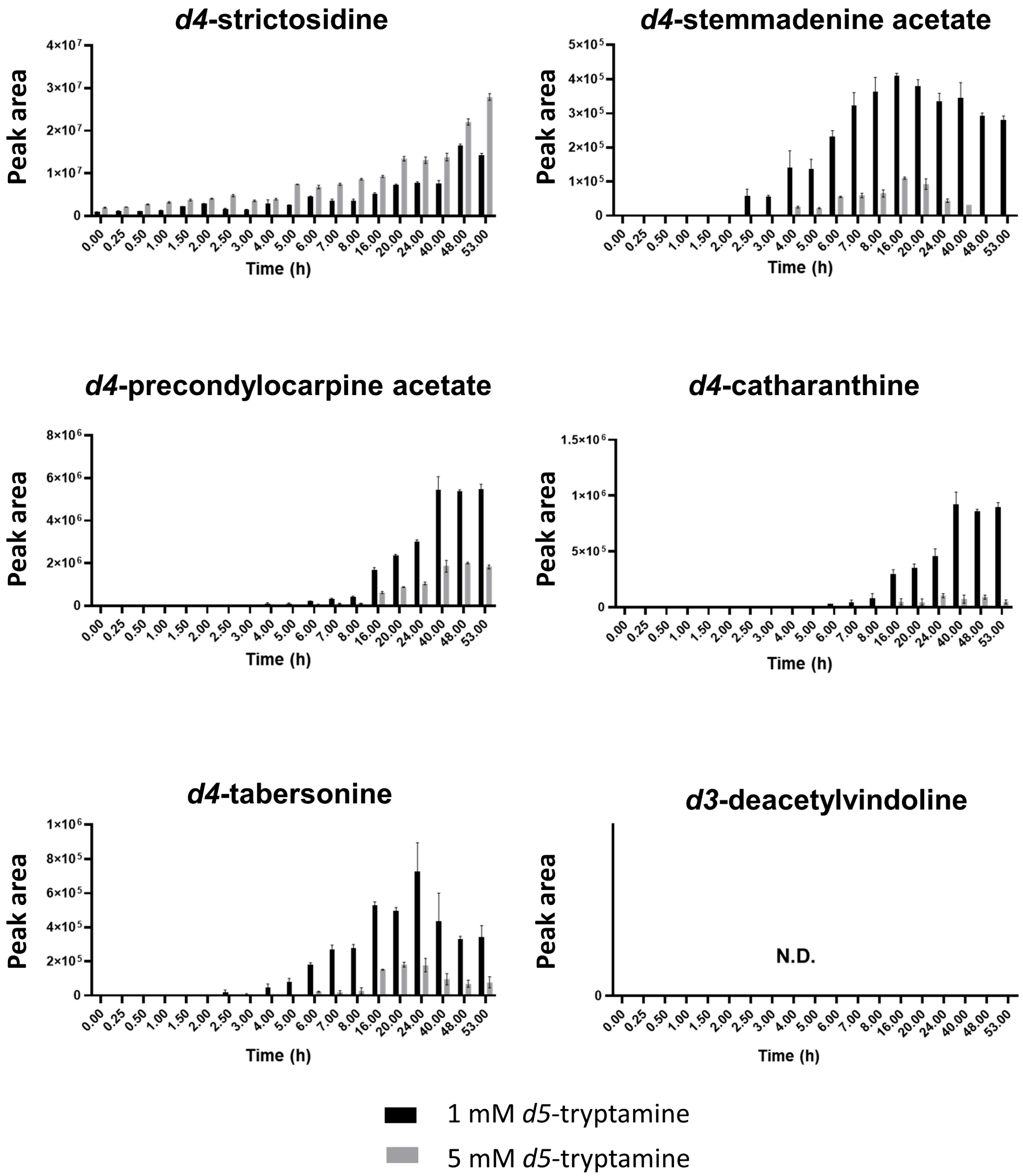


**Figure S2**

Continued on next page →

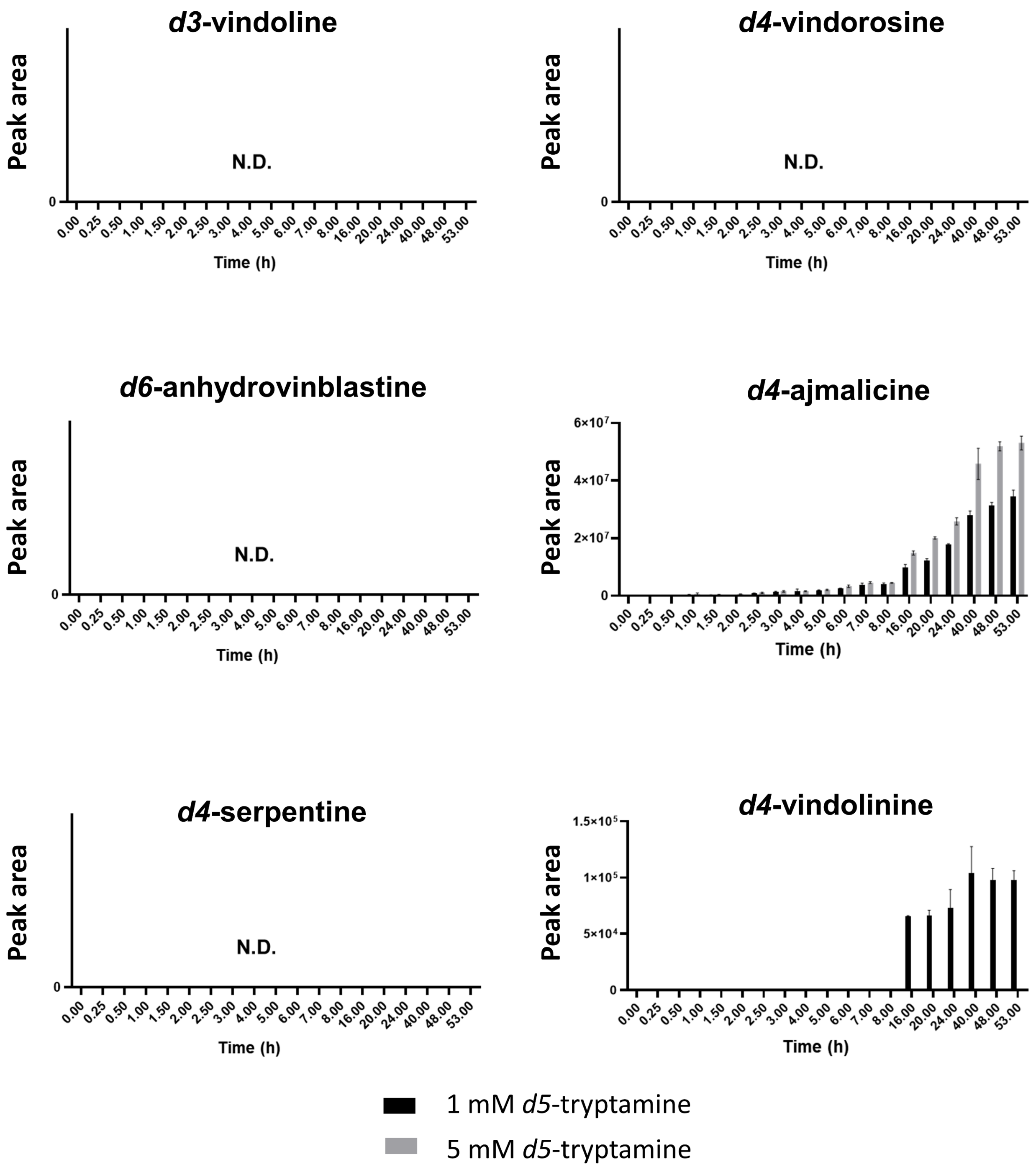


**Figure S2.** Incorporation of *d5*-tryptamine in bulk protoplasts derived from *C. roseus* leaf. An aliquot of protoplasts was taken at each time point, extracted and subjected to LC-MS. Comparison of incorporated products over 53 hours at different concentrations of *d5*-tryptamine (1 mM and 5 mM) is shown.

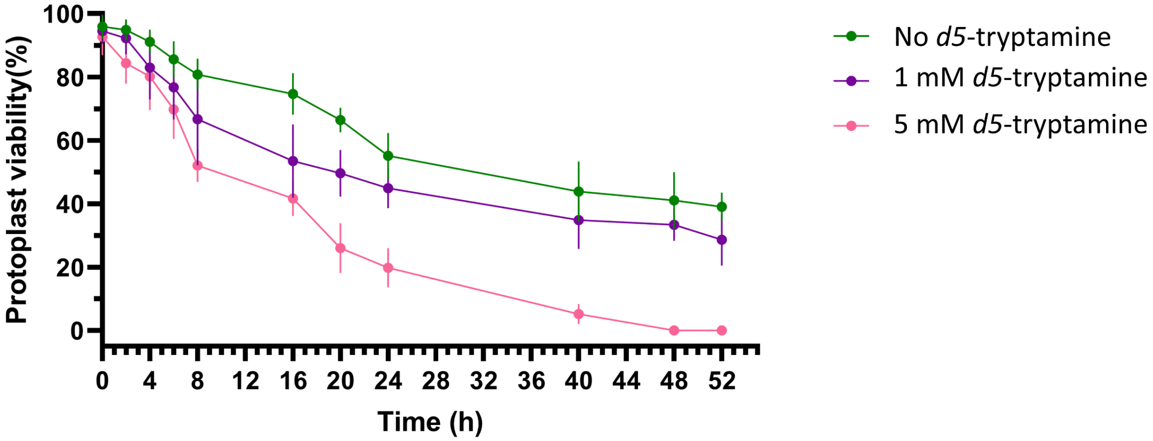


**Figure S3.** Analysis of cell viability under three different feeding conditions: 0 mM, 1 mM, and 5 mM of *d5*-tryptamine in MM Buffer (0.4 M mannitol, 20 mM MES, pH 5.7). Viability of protoplasts was measured by counting viable protoplasts stained by fluoresceine diacetate divided by the total protoplast counted under bright field on a hemocytometer. Counts were performed in triplicate. The means are plotted and error bars indicate standard deviation.

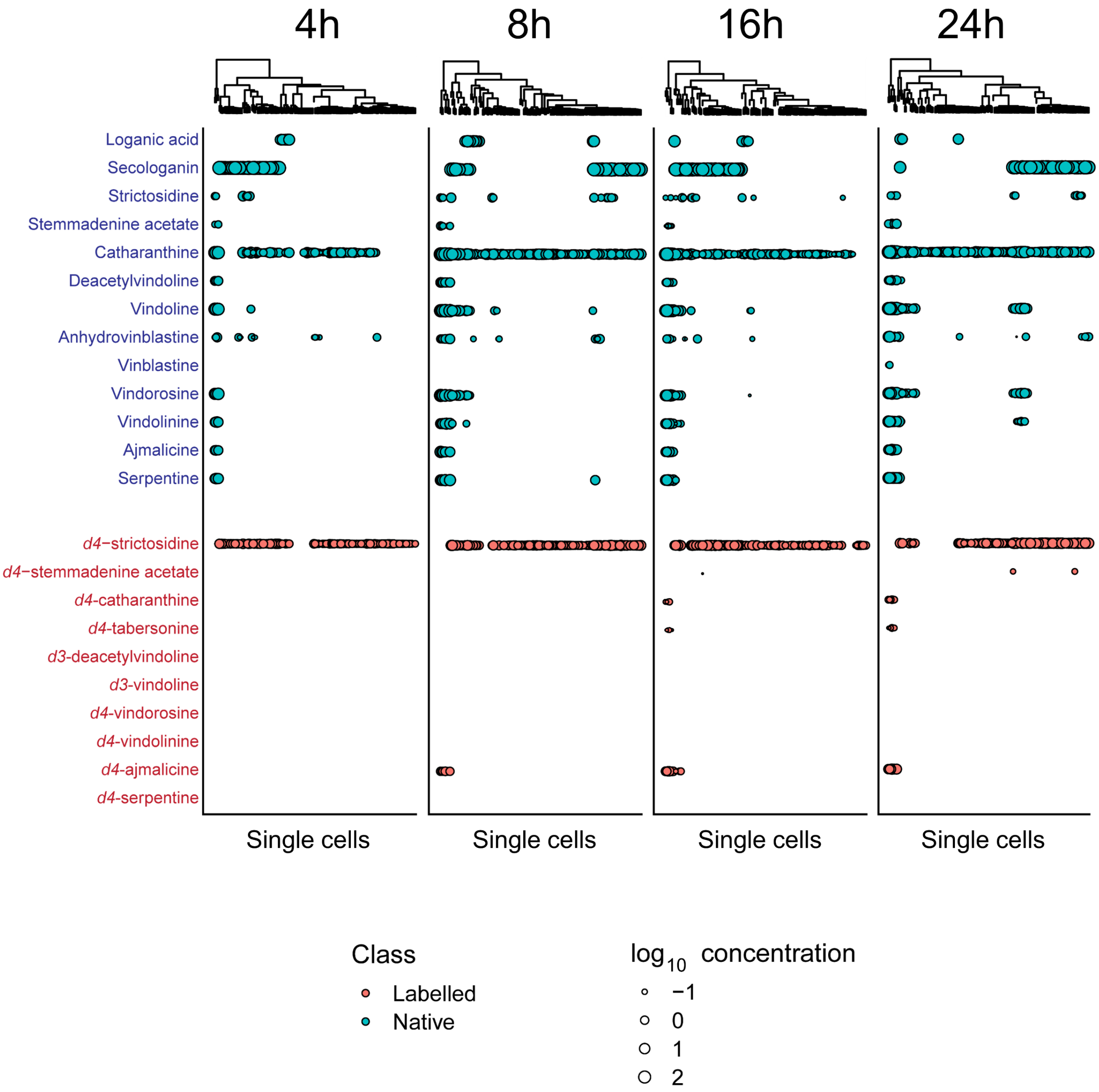


**Figure S4.** Metabolic profile of both labeled and native target compounds quantified in single protoplasts over a 24-hour time course of the protoplast feeding experiments. Protoplasts are grouped by hierarchical clustering. Compounds in blue represent unlabeled metabolites, while compounds in red are derived from *d5*-tryptamine. The size of the dot represents the $\log_{10}$ of concentration (mM) of the metabolites in the corresponding cells.

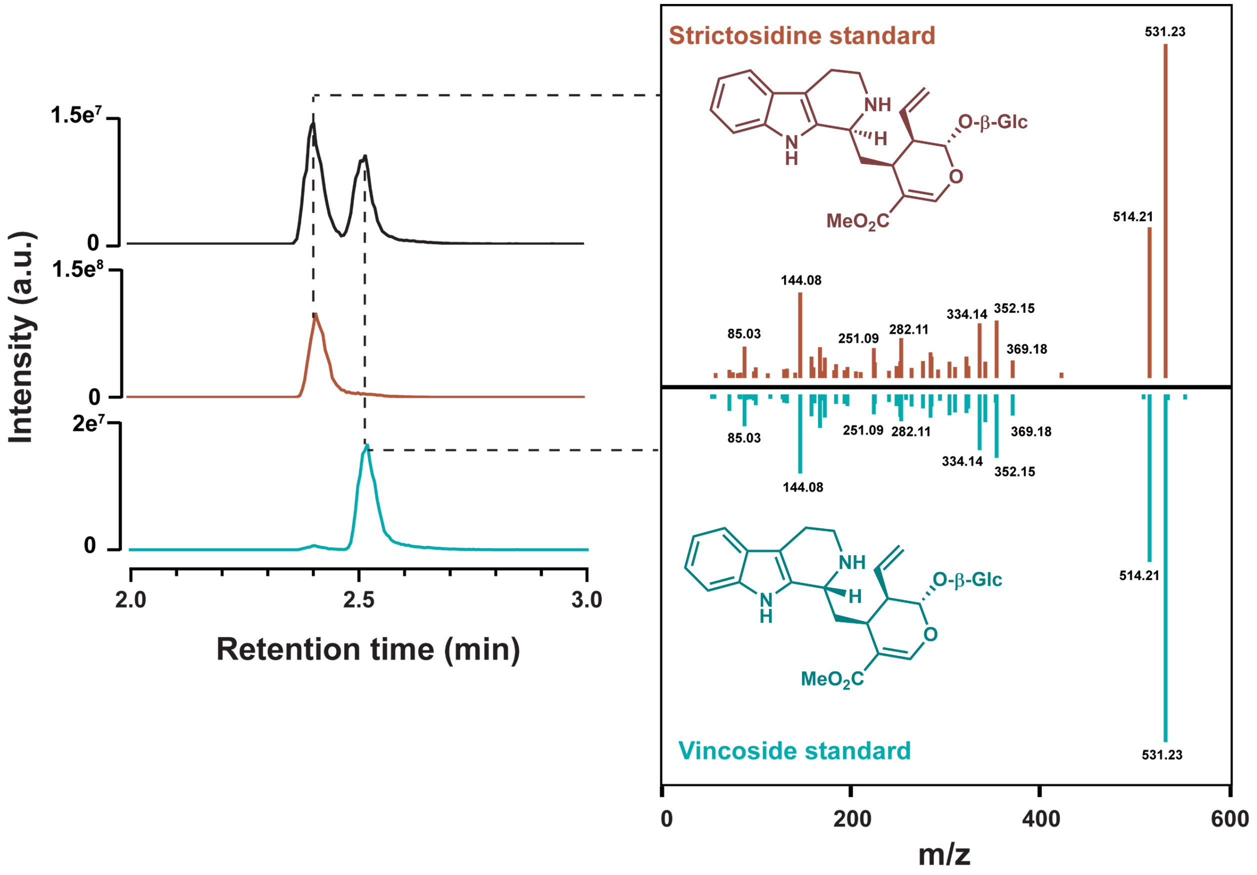


**Figure S5.** Extracted ion chromatograms (EICs) and MS/MS spectra of strictosidine and vincoside standards.

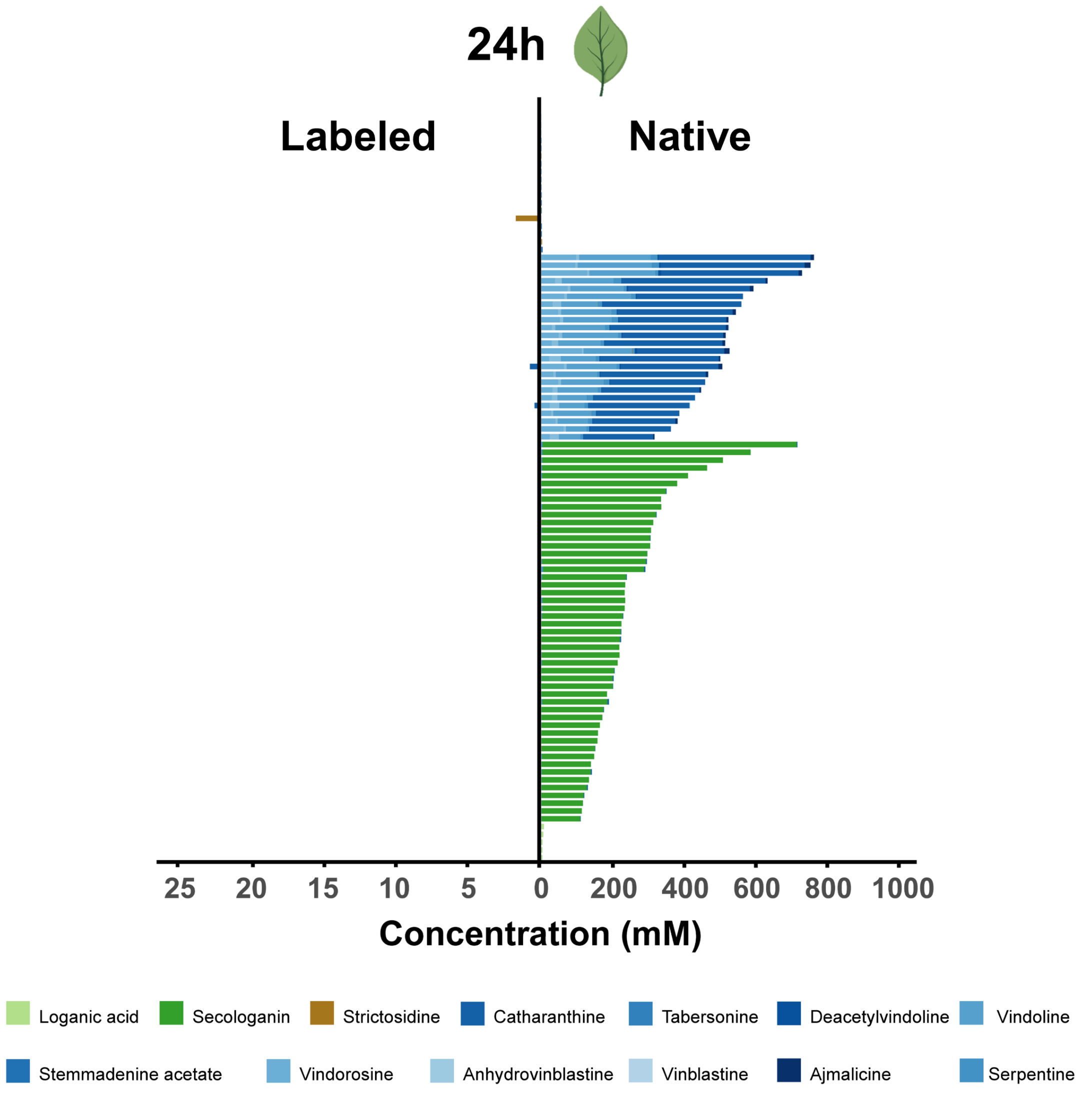


**Figure S6**. Single cell metabolic profile of protoplasts derived from an intact leaf fed through the petiole with 1 mM *d5*-tryptamine for 24 hours. Out of 176 analyzed cells, 96 cells with quantifiable levels of labeled and native target compounds were visualized. Each individual bar represents a single cell. Data are presented as mM concentration.

(a)

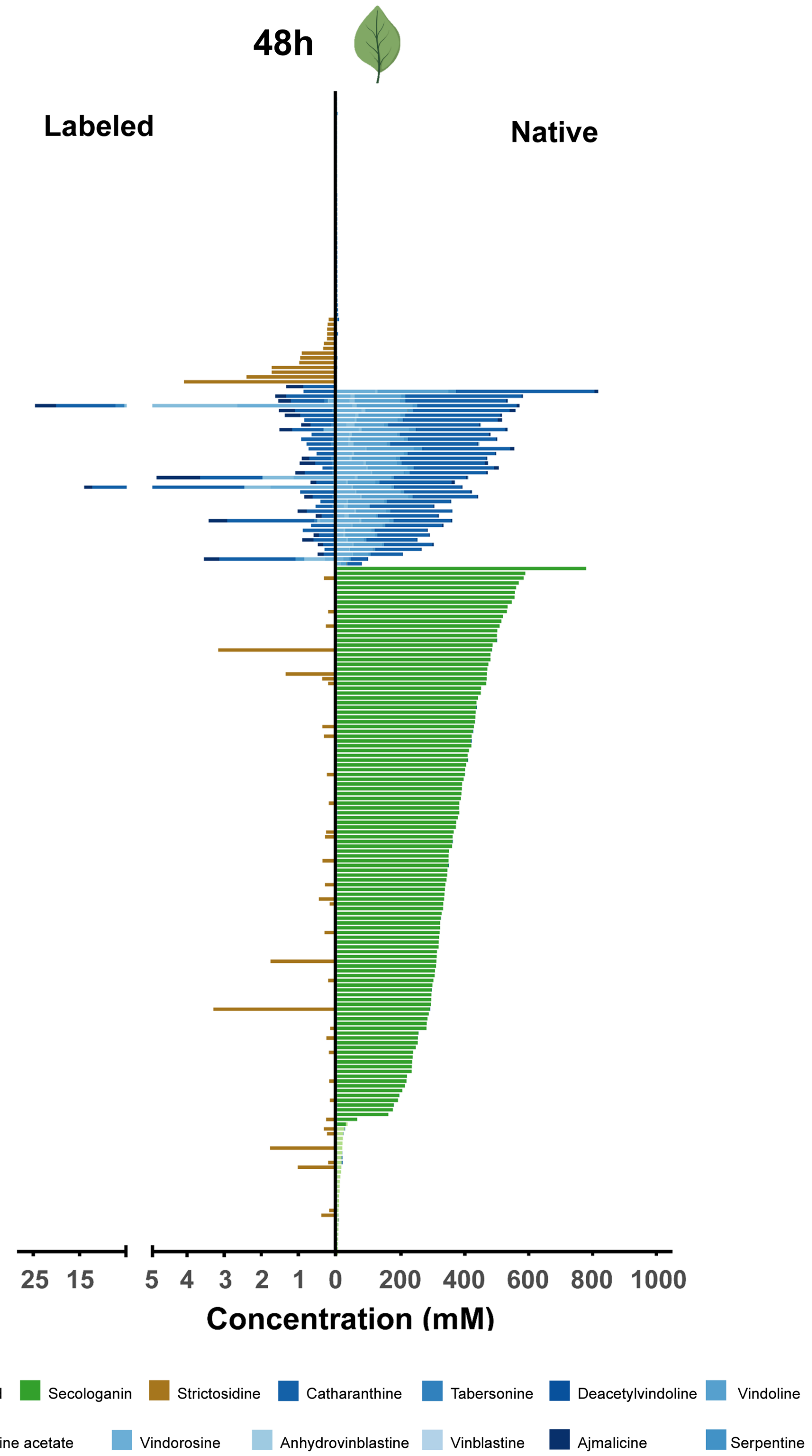
48h
Labeled
Native
25 15
5 4 3 2 1 0 200 400 600 800 1000
Concentration (mM)
Loganic acid
Secologanin
Strictosidine
Catharanthine
Tabersonine
Deacetylvindoline
Vindoline
Stemmadenine acetate
Vindorosine
Anhydrovinblastine
Vinblastine
Ajmalicine
Serpentine

**Figure S7** Continued on next page →

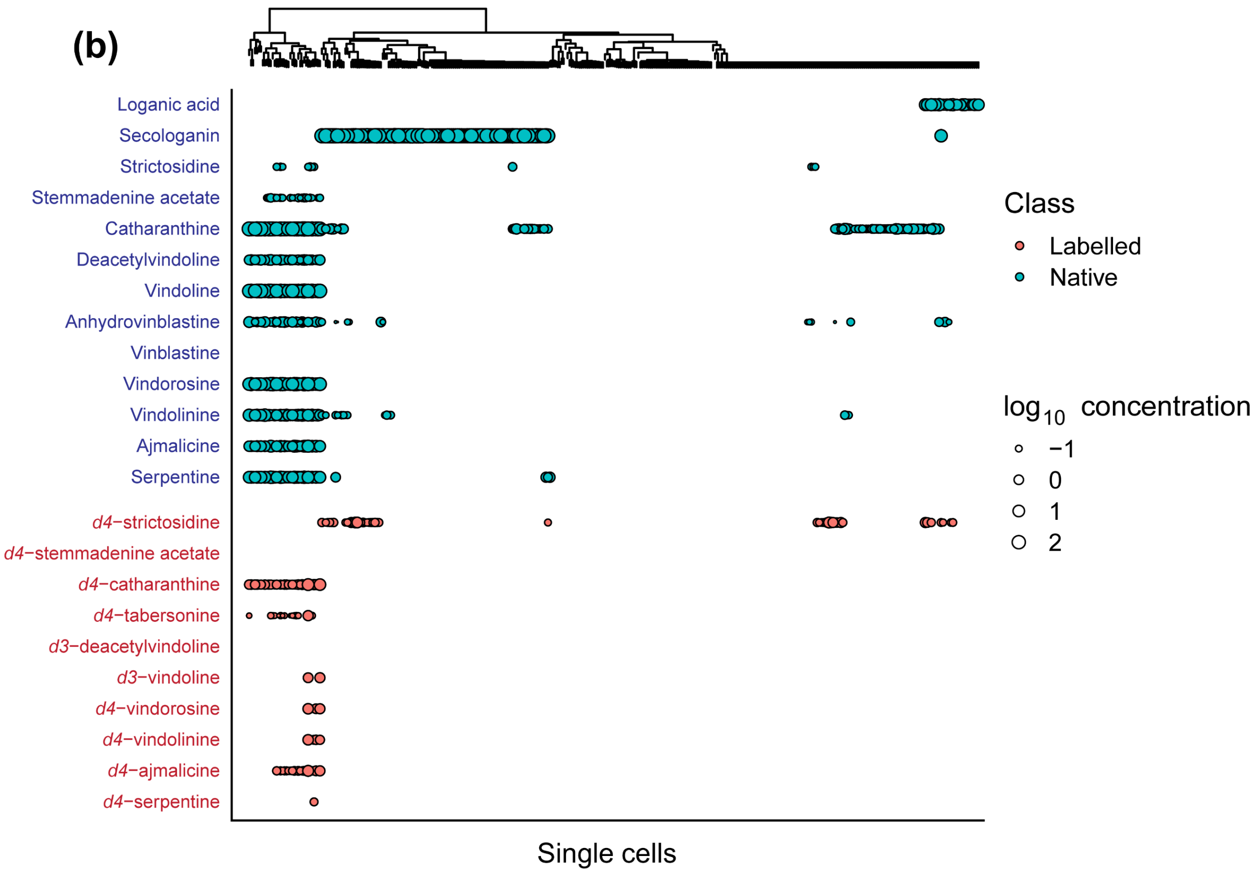


**Figure S7.** Single cell metabolic profile of protoplasts derived from an intact leaf fed through the petiole with 5 mM *d5*-tryptamine for 48 hours. Data are presented as a stacked bar chart **(a)** and a dot plot **(b)**. In the stacked bar chart **(a),** each individual bar represents a single cell and data are presented as mM concentration. In the dot plot **(b),** cells are grouped by hierarchical clustering. Compounds in blue represent unlabeled metabolites, while compounds in red are derived from *d5*-tryptamine. The size of the dot represents the $\log_{10}$ of concentration (mM) of the metabolites in the corresponding cells.

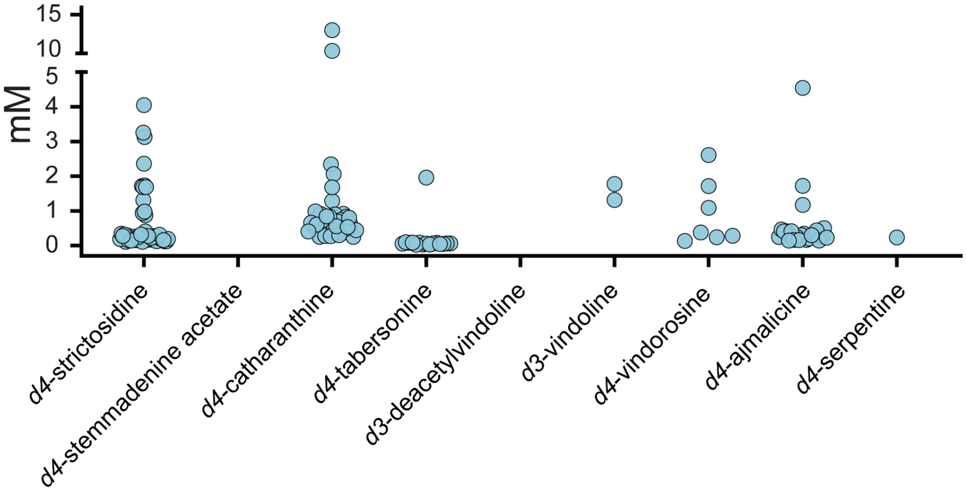


**Fig. S8** Intracellular concentration in mM of labeled products in protoplasts derived from an intact leaf fed through the petiole with 5 mM *d5*-tryptamine for 48 hours (Fig. **1c**). Dots represent the concentration of detected labeled products in individual cells.

**Table S1.** Number of cells analyzed in scMS experiments.

| Experiment | | Number of cells analyzed |
|---|---|---|
| **Feeding protoplast** | 4h | 179 |
| | 8h | 172 |
| | 20h | 158 |
| | 24h | 174 |
| **Feeding intact leaf** | 24h | 176 |
| | 48h | 372 |

**Table S2.** Analytical parameters of the compounds quantified in this study.

| Compound | Calibration range (nM) | Regression equation[a] | Correlation Coefficient ($R^2$) | LOQ[b] (nM) |
|---|---|---|---|---|
| Loganic acid | 2-2000 | y = 12248x + 6150 | 0.9999 | 2 |
| Secologanin | 10-1000 | y = 16165x + 84261 | 0.9988 | 10 |
| Strictosidine | 0.1-500 | y = 855571x + 379613 | 0.9999 | 0.1 |
| Catharanthine | 0.1-400 | y = 1039954x + 283600 | 0.9997 | 0.1 |
| Deacetylvindoline | 0.05-500 | y = 750146x + 1400257 | 0.9985 | 0.05 |
| Vindoline | 0.1-500 | y = 1186763x + 1795458 | 0.9991 | 0.1 |
| Vindorosine | 0.1-500 | y = 846541x + 597335 | 0.9996 | 0.1 |
| Tabersonine | 0.1-500 | y = 167316x + 3653934 | 0.9982 | 0.1 |
| Serpentine | 0.5-500 | y = 1196468x + 269956 | 0.9994 | 0.5 |
| Vindolinine | 0.05-150 | y = 2210449x + 710425 | 0.9993 | 0.05 |
| Anhydrovinblastine | 0.02-50 | y = 1976113x + 64437 | 0.9999 | 0.02 |
| Vinblastine | 0.05-200 | y = 1844895x + 942822 | 0.9994 | 0.05 |
| Stemmadenine acetate | 0.05-50 | y = 7047298x + 1129252 | 0.9993 | 0.05 |
| Ajmalicine | 0.05-200 | y = 1013368x + 535542 | 0.9993 | 0.05 |

[a]Each point of calibration curve was measured in triplicate.

[b]The LOQ was estimated in the lowest analyte concentration injected that yielded a signal-to-noise (S/N) ratio of ≥10 in three replicates.

**Table S3**. Data used to calculate the rate of accumulation of *d4*-strictosidine in the protoplast feeding experiment

| | **Time 4 h** | **Time 8 h** | **Time 20 h** | **Time 24 h** |
|---|---|---|---|---|
| Number of cells analyzed | 179 | 172 | 158 | 174 |
| Secologanin-containing cells (epidermal) | 54 | 58 | 53 | 68 |
| Percentage of epidermal cells | 30 | 34 | 33 | 39 |
| *d4*-strictosidine-containing cells | 154 | 149 | 133 | 123 |
| Total *d4*-strictosidine produced (fmol) | 1130 | 1810 | 3438 | 4371 |
| *d4*-strictosidine/cell produced (fmol) | 20.61 | 31.21 | 64.86 | 64.28 |
| Rate (fmol/h)/cell | 5.15 | 3.90 | 3.24 | 2.68 |
